# Single-Cell Behavioromics Enables Fate Prediction of Individual Hematopoietic Stem Cells

**DOI:** 10.64898/2026.10.06.757119

**Authors:** Alisa B. Chankhunthod, Rhys Sorenson-Graff, Juliane Mercoli, Takashi Ishida, Stacey Dozono, Rachel Wellington, Youyi Fong, Brandon Hadland, Helong Gary Zhao

## Abstract

Achieving near-single-cell precision in the prospective identification of long-term engrafting hematopoietic stem cells (LT-HSCs) remains challenging, likely because current static, point-in-time single-cell phenotyping technologies do not capture the dynamic nature of cell fate decisions. Here, we overcome this barrier by longitudinally tracking individual hematopoietic stem and progenitor cells (HSPCs) in *ex vivo* niche cultures using live imaging, coupled with downstream immunophenotyping, single-cell transcriptomics, and transplant assays. Deep learning analysis of live imaging data achieved 98.5% average precision in predicting serially-engrafting LT-HSC (serial-LT-HSC) identity from each cell’s behaviorome (the totality of observed behaviors), outperforming currently available technologies. Serial-LT-HSCs exhibited lower motility among HSPC populations, which was associated with reduced Activin signaling activity and elevated expression of cell adhesion genes. Correspondingly, antagonizing Activin signaling by Inhibin-A treatment in *ex vivo* niche cultures suppressed HSPC motility while increasing the frequency of post-expansion serial-LT-HSCs, revealing a novel role for Activin signaling in HSPC behavior and fate determination. Collectively, this study establishes a broadly applicable framework that connects behavioromic analysis and single-cell transcriptomics to enable near-single-cell precision in predicting stem cell fate and uncover molecular mechanisms governing behavior-fate relationships.

## Introduction

Hematopoietic stem cell transplantation (HSCT) remains a pillar of modern medicine, offering cures for many blood and immune disorders. The therapeutic efficacy of HSCT depends on rare long-term hematopoietic stem cells (LT-HSCs) capable of durable engraftment, self-renewal, and multilineage reconstitution^1^. Despite their central importance in HSCT and stem cell research, prospective identification of live LT-HSCs at single-cell-level resolution has been challenging. Decades of progress in immunophenotyping has enabled enrichment of rare LT-HSCs from mouse and human hematopoietic cells^2-9^; however, the resulting populations often remain heterogeneous, contaminated with various proportions of progenitors with limited engraftment and self-renewal potential^3,4,10^.

“Snapshot” single-cell genomic profiling technologies, such as single-cell RNA sequencing (scRNA-seq) and single-cell ATAC-seq, have substantially improved our understanding of hematopoietic heterogeneity, revealing transcriptional and epigenetic differences among stem and progenitor compartments^4,8,9,11-14^. However, static molecular features do not fully capture the functional potential or fate of HSPCs, which are shaped by dynamic niche interactions, temporally arranged transactivation programs, and continuous cellular state transitions^12,15,16^.

Live-cell imaging provides a unique means to observe dynamic cellular states that may more precisely predict fate^17^. Such behaviors encode time-resolved information about stemness that is inaccessible by static marker-based analyses or snapshot omics profiling^12^. For example, we previously showed that serially-engrafting LT-HSCs (serial-LT-HSCs) from fetal liver tend to divide symmetrically in a self-renewing fashion, whereas progenitors more readily undergo asymmetric division as they differentiate^4^. By tracking individual cells as they migrate, divide, and interact with their niche in a high-throughput way, we can construct single-cell behavioromes (the totality of observed cellular behaviors) and apply deep learning (DL) methods to identify functional HSPC fates based on their distinctive behavioromic features.

Here we present an approach for prospective prediction of serial-LT-HSC fate by DL analysis of single-cell behavioromes and provide a mechanistic link between behavior and fate.

## Results

### HSPC behavioromic profiling predicts serial-LT-HSC fate

With the goal of accurate fate prediction in live cells, we chose live imaging as a non-invasive and non-destructive approach for identifying dynamic features of HSPCs (Fig. 1A). Leveraging our established methodologies^4^, murine fetal liver (FL) HSPCs (CD45^+^GR1^-^F4/80^-^ SCA1^+^EPCR^+^) were fluorescently labelled and individually index-sorted onto a fetal liver endothelial cell (FL-EC) layer for live imaging of individual HPSCs in their native niche. Using this system, we previously showed that serial-LT-HSCs faithfully maintain a homogenous immunophenotype of CD45^+^GR1^-^F4/80^-^SCA1^+^EPCR^+^ESAM^+^ after a 14-day in-niche expansion^4^ (Suppl. Fig. 1A). Building on this system that links high-throughput live-cell imaging to the functional outcome of individual HSPCs, we performed time-resolved DL analysis of the live imaging data during the first 60 hours of HSPC + FL-EC co-culture^4^, aiming to extract behavioromic features and map them to fate / engraftment outcome (Fig. 1B). Specifically, we trained a DL classification model using 80% of a dataset of three independent experiments (N=288 co-cultures) and validated the model’s performance with the 20% held-out data. The serial transplantation assay outcome of each seeding cell, as inferred by post-expansion immunophenotype, was used as the ground-truth label of live imaging data (“serial-LT-HSC” vs. “other HSPC”, with a naturally occurring ratio of 1:0.6 ∼ 1:1.5 per experiment). When tested on different held-out data, the trained DL model achieved an average precision of 0.967 ∼ 0.985 (area under the precision-recall curve) in identifying serial-LT-HSCs based solely on their behavioromic features (Fig. 1C, D). These findings indicate that single-cell behavioromic features deduced from live-cell imaging data are sufficient to accurately distinguish serial-LT-HSCs from other HSPCs.

**Figure 1.**
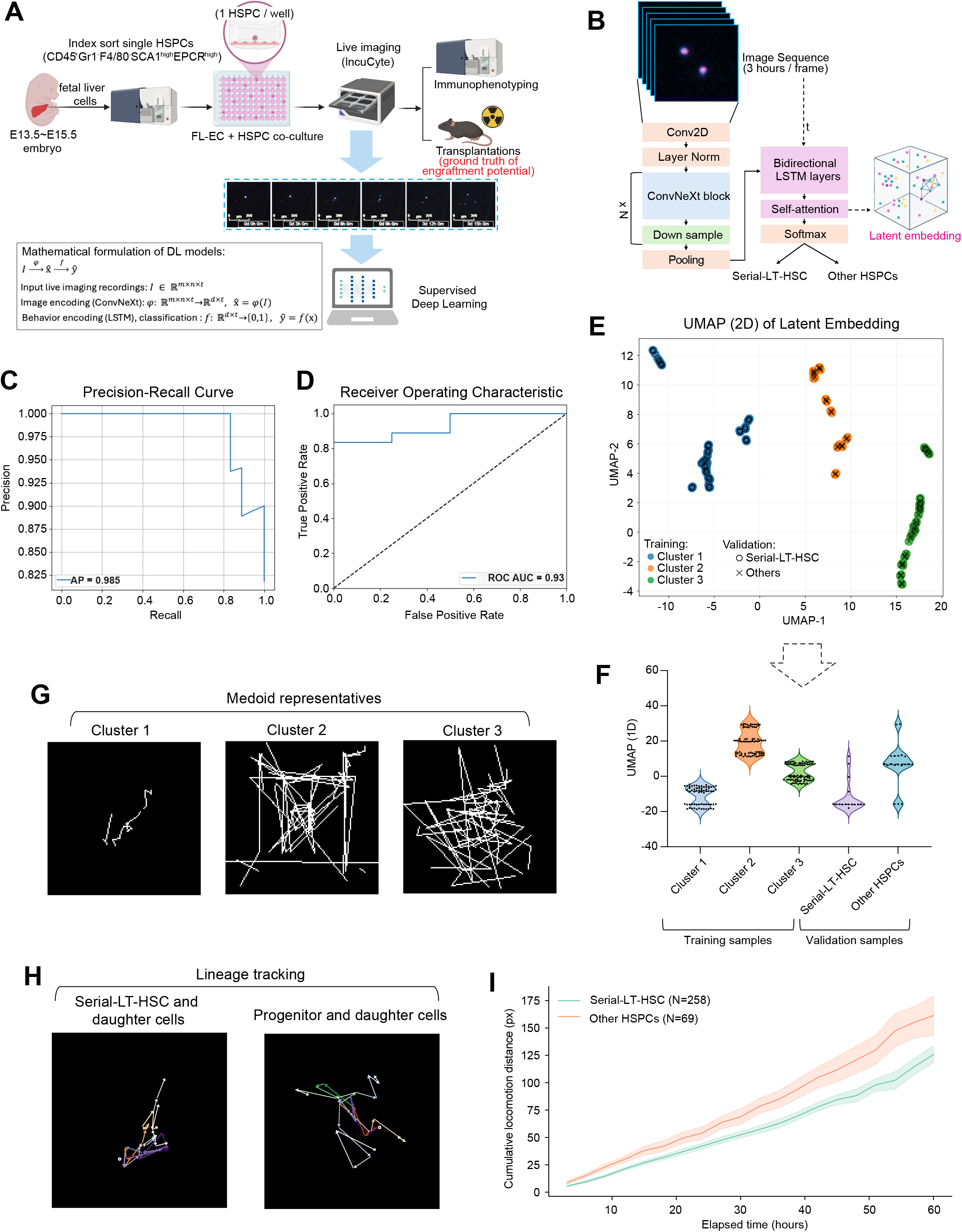
HSPC behavioromic profile correlates with cell engraftment potential. **(A)** Overview of the study design. The keystone of the experimental platform is the linkage among behaviorome, immunophenotype, and engraftment outcome ground truth. **(B)** DL model architecture (see methods for detailed description). **(C-D)** Precision-Recall curve and ROC (Receiver Operating Characteristic) curve of the DL model trained on a dataset containing two independent experiments. The performance was assessed on left-out validation samples. AP: average precision. AUC: area under curve. **(E)** 2D UMAP projections of the latent embeddings generated by trained DL models using live imaging data. **(F)** Violin plot of the 1D UMAP latent embeddings among clusters and cell types in the training and validation samples. The y axis represents the samples’ coordinates in latent space. **(G)** Representative cell-locomotion traces, selected as the cluster medoids (central data point of a cluster) in the UMAP projection. **(H)** Representative lineage-tracking images of a serial-LT-HSC and other HSPC. Cells in each well are color-coded, and arrows indicate cell movements. **(I)** Cumulative locomotion distance of index-sorted serial-LT-HSC or other HSPC and their progenies in co-culture, pooled from 4 independent experiments. Lines indicate the geometric mean for each group and shaded regions represent 95% confidence intervals.

### Serial-LT-HSCs exhibit low motility in the endothelial niche

To stratify FL HSPCs by their behavioromes, we extracted the latent embeddings of cell behaviors from the DL model’s self-attention layer output and visualized them using UMAP (Uniform Manifold Approximation and Projection). 2-dimensional UMAP identified 3 distinct clusters of HSPCs based on their behavioral patterns (Fig. 1E). To compare the distribution of each cell type’s behavior, the latent embeddings were further projected onto 1-dimensional UMAP, retaining three clusters on the linear scale (Fig. 1E, F). Serial-LT-HSCs were enriched in Cluster 1, while other HSPCs were enriched in Cluster 2 and 3 (Fig. 1F), representing distinct behavioromic patterns. Cells in Cluster 1 demonstrated less stochastic locomotion as shown by cell traces over the duration of live imaging (Fig. 1F-G, Suppl. Fig. 1B-C). With hierarchical cell lineage tracking, we observed that daughter cells of serial-LT-HSCs tended to retain low motility while daughter cells of other HSPCs resembled their parental cells as being highly mobile (Fig. 1H-I).

### The low motility of serial-LT-HSC is associated with low niche-derived Activin signaling and high expression of cell adhesion genes

To uncover the molecular mechanism governing the behavioromic features of HSPCs in the FL-EC niche, we analyzed scRNA-seq data from both HSPCs and FL-EC^4^. We hypothesized that HSPC motility is directly regulated by niche-to-HSPC signaling via cytokines or surface ligands (Fig. 2A). Cell-cell communication (CCC) analysis of ligand-receptor pairs between FL-EC and HSPCs by NicheNet^18^ revealed that serial-LT-HSCs had reduced specific interaction potentials between Activin ligands and receptors compared to other HSPCs, when focused on the genes differentially expressed between serial-LT-HSCs and other HSPCs (Fig. 2B-C). This was validated using CellPhoneDB^19^, an orthogonal CCC inference method (Fig. 2D). As Activin signaling regulates cell migration programs^20^, we postulated that the lower Activin signaling in serial-LT-HSCs may contribute to their lower motility. Supporting this, compared to other HSPCs, serial-LT-HSCs showed increased expression of genes associated with cell adhesion, notably including many target genes (e.g. *Tgfbi, Tln*, and *Smad7*) of TGF-β signaling^21,22^ (Fig. 2E), in addition to other previously described features of fetal/neonatal LT-HSCs such as increased vesicular trafficking^23^ (Fig. 2E), enhanced interferon response^24,25^ (Fig. 2E), and reduced protein translation and metabolic activity^4^ (Suppl. Fig. 2A). Correspondingly, serial-LT-HSCs had higher gene set scores for canonical TGF-β signaling and cell adhesion, and lower gene set scores for Activin signaling (Fig. 2F). In aggregate, these data indicate that the lower motility of serial-LT-HSCs relative to other HSPCs is associated with a reciprocal balance between Activin and TGF-β signaling, which may promote expression of downstream proteins that strengthen adhesion to and retention within the FL-EC niche.

**Figure 2.**
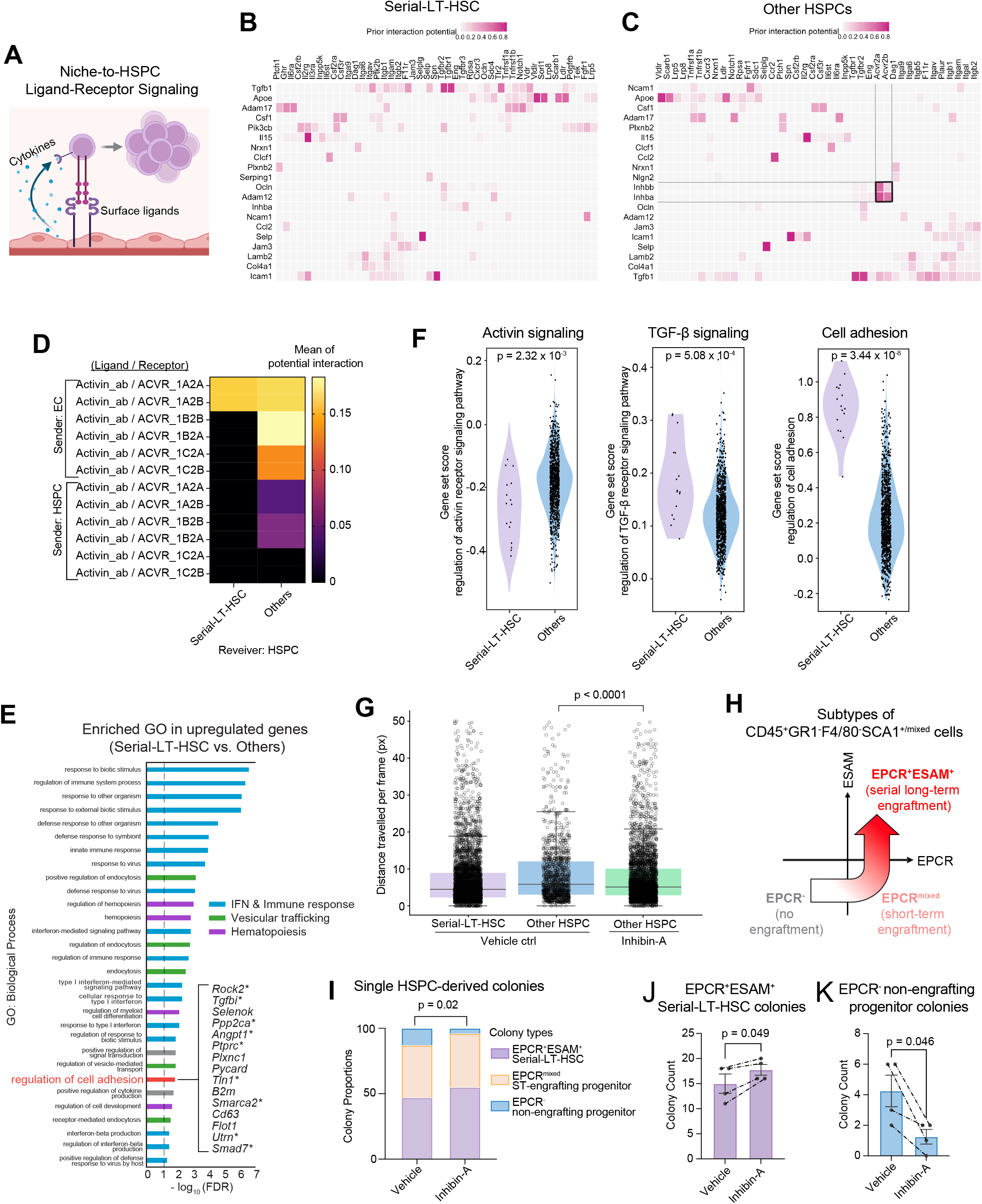
Niche-derived Activin signaling influences cell behaviors and distinguishes serial-LT-HSCs from other HSPCs. **(A)** Diagram illustrating the two types of ligand-receptor interactions between the FL-EC niche and HSPCs examined using CCC analysis from scRNA-seq data. **(B-C)** Ligand-receptor interactions from niche FL-ECs to serial-LT-HSCs and other HSPCs, inferred by NicheNet. Activin-to-Activin Receptor interactions unique to other HSPCs are marked by bold boundary lines. **(D)** Ligand-receptor signaling to serial-LT-HSCs and other HSPCs, inferred by CellPhoneDB. **(E)** Significantly enriched gene ontology (GO) terms (biological process) among upregulated genes in serial-LT-HSCs compared to other HSPCs. GOs are color-coded by their functional categories. Upregulated genes present in the gene set of “regulation of cell adhesion” are shown; *indicates target genes of canonical TGF-β-Smad signaling pathway. IFN: interferon. **(F)** Gene set scores for Activin signaling (GO:0032926, 0032927), TGF-β signaling (GO:0030511), and cell adhesion (GO:0030155) from scRNA-seq data of post-expansion serial-LT-HSCs (N=16) and other HSPCs (N=1014). P values were determined by Mann-Whitney U test. **(G)** Velocity of cell movements (averaged per frame) of index-sorted HSPCs and their progenies treated with Inhibin-A or vehicle (control). Serial-LT-HSC: N=4618 mother-daughter cell lineage, Other HSPC with vehicle ctrl: N=1128 mother-daughter cell lineage, and HSPC with Inhibin-A treatment: N=3977 mother-daughter cell lineage. Statistics were performed by Mann-Whitney U test with Holm correction. **(H)** Diagram illustrating the different engraftment outcomes inferred by post-expansion colony immunophenotypes^4^. Flow cytometry gating of the colonies is shown in Supplementary Figure 2B. ST: short-term. **(I)** Proportions of post-expansion colony types detected following co-culture of individual index-sorted HSPCs treated with Inhibin-A or vehicle (control). P value determined by Chi-square contingency test, from 4 pooled independent experiments. **(J-K)** Total number of serial-LT-HSC (J) and non-engrafting progenitor (K) colonies following co-culture of individual index-sorted HSPCs treated with Inhibin-A or vehicle (control) (N=48 HSPCs per experiment). P values were determined by paired t test, pooled from 4 independent experiments. Dashed lines indicated paired data collected from each separate experiment.

### Antagonizing Activin signaling with Inhibin-A reduces HSPC motility and enhances serial-LT-HSC fate during FL-EC co-culture

To investigate if altering Activin signaling affects the motility of HSPCs observed in the FL-EC niche, we repeated co-culture experiments in the presence of recombinant Inhibin-A, an endogenous antagonist of Activin signaling. Live imaging and cell tracing showed that Inhibin-A treatment reduced the motility of other HSPCs in the FL-EC niche, compared to vehicle control, closer to the baseline level of serial-LT-HSCs (Fig. 2G). We also enquired whether the observed effect on HSPC behavior by Inhibin-A treatment impacted their stemness. Indeed, we observed a modest but significant increase in the frequency of serial-LT-HSC colony type across four independent experiments (Fig. 2H-J, Suppl. Fig. 1A). When we further subdivided other HSPCs into short-term (ST)-engrafting and non-engrafting progenitor categories based on their post-expansion immunophenotypes, as previously described^4^ (Fig. 2H, Suppl. Fig. 2B), there was a corresponding significant reduction in the non-engrafting progenitor subtype (Fig. 2K). Together, these results show that antagonizing Activin signaling with Inhibin-A alters HSPC motility coupled with a shift toward enhanced stem cell potential.

## Discussion

The ability to accurately identify and isolate functional HSCs, particularly serially-engrafting LT-HSCs, is fundamental to both the study of HSC biology and the development of HSC-based therapies. Decades of immunophenotypic refinement have yielded substantial enrichment of LT-HSCs from murine and human hematopoietic tissues^3,5-9^. Recent advances in single-cell transcriptomic and epigenomic profiling have further illuminated the heterogeneity within immunophenotypically defined HSC populations^4,8,9,11-14^. Yet prospective identification of live LT-HSCs with near-single-cell precision has remained challenging. Here, we overcome this barrier using live-cell imaging integrated with DL to accurately link HSPC behavioromics to serial-LT-HSC cell fate.

Prior studies have typically investigated specific behaviors of HSCs (*e*.*g*. chemotaxis^26^) motivated by molecular mechanistic discoveries (*e*.*g*. critical role of CXCL12-CXCR4 signaling^27^). Our study, however, adopted a comprehensive behavioromic approach, capturing a large variety of HSPC behaviors during niche interactions and analyzing the behaviorome as a whole by DL. Through embedding and classifying behavioromic profiles in high-dimensional latent space, we identified time-resolved patterns that correlate precisely with cell fate. Lower motility was a prominent feature associated with serial-LT-HSCs, representing one of likely many more subtle dynamic behavioral features^4^ distinguishing serial-LT-HSCs from other HSPCs.

Our multi-omics framework further links observable behavioromic features to niche-derived signaling pathways and associated cell fates, extending a paradigm of niche-dependent fate decisions in hematopoiesis^4^. We identified Activin signaling as one of several pathways regulating the broader HSPC behaviorome and uncovered reciprocal changes in Activin and TGF-β signaling activity between serial-LT-HSCs and other HSPCs. This finding is consistent with evidence from human cell lines and in mouse mesenchymal stem cells showing that TGF- β and Activin pathways can exert mutually antagonistic effects through differences in receptor usage, SMAD complex formation and feedback regulation, and noncanonical downstream signaling^28,29^. Given the critical role of TGF-β in promoting HSC quiescence and retention in the bone marrow niche^21,22^, our findings suggest that the reciprocal interactions between Activin and TGF-β pathways may fine tune expression of cell-adhesion proteins contributing to the balance between niche retention/adhesion and migration/mobilization.

In addition to modulating cell motility, we further observed a modest, though statistically significant, effect of Activin inhibition on HSPC fate, consistent with a model in which crosstalk among multiple signals collectively shapes serial-LT-HSC state transitions. This finding suggests that integrating additional behavior-associated pathways could enable more effective manipulation of stem cell fates. More broadly, this framework could be applied to predict and engineer the fates of human HSCs and other stem cells for therapeutic applications across diverse developmental and tissue-specific contexts by manipulating niche-instructed behavior-fate relationships.

Although our *ex vivo* platform and DL framework enables correlative studies linking live cell behaviors, niche pathways, and stem cell fates, it does not fully recapitulate the complexities of the *in vivo* HSPC niche, including the influence of non-EC niche populations, such as mesenchymal stromal cells^30^. Additionally, in the present study, the behavioromics and transcriptomics analyses were not fully integrated into a shared latent representation that directly maps specific behaviors to predicted gene regulatory pathways. Future work will address these limitations by extending behavioromic analysis to more complex hematopoietic organoid systems using high-content 3-dimensional confocal live imaging devices and by developing explainable latent variable models that integrate behavioromics with molecular multi-omics.

## STAR Methods

### Dissection of embryos and cell sorting

E13.5-E15.5 embryos were harvested from pregnant C57BL6/J females mated with C57BL6/J males and washed with PBS with 2% FBS. Embryo age was determined by the date of observed maternal plugging and further confirmed by previously described morphologic criteria^4^. Fetal livers were dissected, pooled in 10 mL conical tubes, and dissociated to a single cell suspension in PBS with 2% FBS by vigorous pipetting followed by passage through a 70um strainer. Samples were then washed in PBS and resuspended in red blood cell lysis buffer (150 mM Ammonium chloride, 10 mM Potassium bicarbonate, 0.1 mM EDTA, pH: 7.2– 7.4) for 5 min at room temperature. Cells were washed once, incubated with anti-mouse CD16/32 (FcRII blocker, BD Biosciences Cat#553141), and stained with the following monoclonal anti-mouse antibodies: GR1 (clone RB6-8C5, eBioscience), F4/80 (clone BM8, Biolegend), CD201/EPCR (clone eBio1560, eBioscience), CD45 (clone 30-F11, Invitrogen), SCA1 (clone D7, eBioscience), ESAM (clone 1GB/ESAM, Biolegend). DAPI (Millipore, Cat#268298) was used to exclude dead cells. In some experiments, FL cells were also stained with CellTrace Far Red (ThermoFisher, Cat#C34564) and LysoBrite (Stem Cell Technologies, Cat# 100-0850) following the manufacturer’s protocol. CD45^+^GR1^-^F4/80^-^SCA1^+^EPCR^+^ cells were sorted out as immunophenotypic FL HSPCs (enriched in functional LT-HSCs, as previously described^4^) and deposited onto FL-EC layers at 1 cell per well in a 96-well plate, by either BD FACSymphony6 or Aria II equipped with BD FACS Diva Software with index sorting capability (Becton Dickinson), in single cell mode with maximum purity mask settings to minimize contaminating cells.

### FL-HSPC and FL-EC co-culture

For co-culture experiments, FL-AKT-ECs^4^ at passage 12 or less were plated at a density of 1 × 10^4^ cells per well on 96-well plates 1–2 days prior to use. Prior to co-culture, FL-AKT-EC layers were washed with serum-free X-VIVO 20 media (Lonza). C57BL6/J (CD45.2) FL-HSPCs were individually index sorted as described above to each well of gelatin-coated (MilliporeSigma, Cat No. ES006B), 96-well, coverglass-bottom, black plates (Corning) containing FL-AKT-EC in serum-free X-VIVO 20 media plus recombinant cytokines (PeproTech): murine stem cell factor (SCF) at 100 ng/mL and thrombopoietin (TPO) at 20 ng/mL.

### Flow cytometric analysis of colonies

Following 14 days of co-culture, 50% of the progeny of each index-sorted FL-HSPC in each 96-well was harvested by gentle resuspension for analysis by flow cytometry. Cells were spun down and re-suspended in PBS with 2% FBS, incubated with anti-mouse CD16/CD32 (FcRII block) and then stained with the following anti-mouse monoclonal antibodies: GR1 (clone RB6-8C5, BD Biosciences), F4/80 (clone BM8, Biolegend), CD201/EPCR (clone eBio1560, eBioscience), CD45 (clone 30-F11, Invitrogen), SCA1 PC (clone D7, eBioscience), CD144/VE-Cadherin (clone eBioBV13, eBioscience), ESAM (clone 1GB/ESAM, Biolegend). DAPI was used to exclude dead cells. Cells were analyzed by either BD FACS Canto II or BD FACS LSR Fortessa equipped with BD FACS Diva Software (Becton Dickinson) and further analyzed by Flow Jo ver.10 software.

A colony was defined as having greater than 3 viable CD45^+^GR1^-^F4/80^-^ cells detected by flow cytometry. Based on our previously published criteria^4^, colonies in which greater than 80% of viable CD45^+^ cells retained GR1^-^F4/80^−^SCA1^+^EPCR^+^ESAM^+^ immunophenotype consistently achieved multilineage engraftment following primary and secondary transplantations in irradiated congenic strain mice and were thus classified as serial-LT-HSC colonies. Colonies with CD45^+^GR1^-^F4/80^−^SCA1^+/mixed^EPCR^mixed^ESAM^-^ immunophenotype established short-term engraftment in primary recipients but could not be serially transplanted and were classified as ST-engrafting progenitors. Colonies with CD45^+^GR1^-^F4/80^−^SCA^mixed^EPCR^-^ immunophenotype did not establish engraftments in primary recipients and were classified as non-engrafting progenitors.

### Live imaging

Live-cell imaging of single index-sorted cells was carried out on the Incucyte SX5 incubator microscope system, in which 96-well plates with FL-EC / HSPC co-cultures were maintained at 37 degrees Celsius and in 5% CO_2_. To distinguish HSPCs from FL-ECs in the co-culture, HSPCs were stained with fluorescent dyes, Celltrace-Far Red and LysoBrite. Celltrace-Far Red labels total protein content (free amine) in live cells, which presents cellular morphology. LysoBrite labels lysosomes (acidic organelles), whose activity and distribution have been associated with HSC functionality^23,31,32^. Brightfield and fluorescent images were captured every 2-3 h during the initial incubation period (48–72 h). Images were captured with a 4X objective lens and digitally stitched into whole-well images for all wells of the 96-well plate. Only fluorescent channels were used in deep learning, whose raw image data were exported as tiff files after filtering out low-intensity background noise signals.

### Image processing and deep learning

Before DL, we cropped raw whole-well images to retain only the minimal areas that contain the trace of dividing and moving cells. Due to the ∼1000-fold size difference between whole-well images (6.4 mm) and mouse HSPCs (<10 μm), previously published cell segmentation tools failed to identify single cells in our images. Therefore, we built a custom program to identify HSPCs and crop out the minimal areas used as the input for DL. The program first identifies cell-like objects with overlayed red (LysoBrite) and blue (Celltrace Far Red) pixels, and then selects target areas by comparing the traces of all cell-like objects over time, assuming non-cell objects do not divide. Details are provided in the python code deposited on Github (https://github.com/FredHutch/Chankhunthod_Sorenson-Graff_et_al_2026).

The dataset of cropped image sequences (blue and red fluorescent channels) were augmented to 6 times in size by random rotations, flips, contrast alterations, and brightness alterations, before being fed into the deep learning pipeline. The DL model was composed of Conv2D, ConvNeXt blocks, bidirectional LSTM layers, self-attention layer, and softmax output labelled to two classes of ground-truth: serial-LT-HSCs and other HSPCs (as determined by post-expansion colony immunophenotype). Image sequence datasets containing independent experiments were split into training and testing sets and only training sets were fed into models with ground-truth labels. Training was terminated when there was no longer reduction of loss over time, at 60 epochs. Model performance was evaluated by precision-recall curve and ROC curve. Details are provided in the python code deposited on Github (https://github.com/FredHutch/Chankhunthod_Sorenson-Graff_et_al_2026).

For UMAP projection and clustering, latent embeddings of cell behavioromes were extracted from the self-attention layer and single cells were projected onto 2-dimensional or 1-dimensional UMAP plots.

### Cell detection in perimetric analysis

Cells were detected per-well and per-frame using the live image fluorescent channels, converted to a grayscale floating-point array of pixels. Before detection, images were normalized in two passes: first, pixels in each frame were equalized so that the mean of its pixels at or below the 75th percentile matched the well-wide average; second, each pixel was zero-centered by subtracting the mean of its dimmest 50% of values across frames.

Cells were detected independently in each frame. First, local maxima were identified using 8-connectivity and split into dim and bright populations by k-means clustering (k = 2) on the intensity of the maxima, fit separately per frame. Bright maxima were used as candidate cells while dim maxima were discarded as background. Blobs surrounding the maxima were defined as connected regions (4-connectivity) of pixels exceeding the Tukey upper fence (Q_3_ + 1.5 × IQR) of all post-normalization pixel values of the frame. Bright maxima falling within the same blob were collapsed to the unweighted centroid of the blob’s pixels; isolated maxima were unchanged. Finally, if the number of candidate cells in a frame exceeded 2.0 × the running maximum count from all preceding frames (initialized at 1.0), that frame and all subsequent frames were removed.

### Mother-daughter cell lineage tracking in perimetric analysis

Mother-daughter cell relationships were assigned and tracked between adjacent frames by optimal assignment (Hungarian algorithm), minimizing total Euclidean distance travelled. To analyze a lineage’s cumulative motility (including prior to division), tracking used a fixed number of slots rather than a variable number of candidate cells: the largest candidate cell count observed in any retained frame of a well-defined the slot count for all frames. For frames with fewer candidate cells, multiple slots shared the parent cell’s coordinate. Assignment proceeded backward in time so that division appeared as a many-to-one merge of daughters-to-parent. A division was recorded at frame t+1 when two or more slots sharing a coordinate at frame t originated from more than one distinct coordinate at t+1. Because an undivided parent is represented by as many slots, shared slots were collapsed to a single lineage; slots diverging at a division were retained separately.

Incorrect assignments and assignments to false positives were removed in four ways. First, links over 50 px were penalized to prioritize shorter assignments; where none existed the link was cut, and that lineage’s frames before and after were treated as separate lineages. Second, lineages were required to contain at least one division during the retained frames, as non-cellular particles that did not divide and could not be confirmed as live cells. Third, because real cells dim over time while non-cellular particles do not, mean intensity of each candidate cell’s blob was regressed on frame index across all frames of dividing lineages by ordinary least squares (polynomial degree = 2). Candidate cells above the fitted curve (internally studentized residual > +2.0) were removed as failing to dim with the cell population. Fourth, lineages observed in fewer than three frames were excluded.

Lineages passing all filters were compared between conditions by cumulative distance, summed from the start of imaging and sampled every 3 h to 60 h. Two conditions were judged to have different motilities if the 95% confidence intervals of their geometric means did not overlap.

### Single cell transcriptomic analysis

scRNA-seq data of HSPCs (NCBI GEO Accession GSM7400740) in co-culture with FL-ECs (NCBI GEO Accession GSM7400741) were loaded and processed in Python using Anndata and Scanpy^33^. serial-LT-HSCs and other HSPCs were annotated per immunophenotyping and serial transplant outcome described previously^4^.

#### Differential expression analysis

Anndata object of HSPCs was normalized by total gene counts (1e4) and log1p transformed. Differential expression genes between serial-LT-HSCs and other HSPCs were analyzed by scanpy.rank_genes_groups with Wilcoxon test.

#### Gene ontology (GO) enrichment analysis

Biological Process GO enrichment was carried out by Toppgene API^34^, with q value by Bonferroni correction as statistical test.

#### Gene set score

gene set score was calculated using scanpy.score_genes with gene sets of specified GOs. Two-group comparison of gene set scores was performed by Mann-Whitney U Test using Scipy.stats.

#### Cell-cell communication (CCC) analysis

*CCC was performed using NicheNet*^18^ and *CellPhoneDB*^*19*^ tools. Nichenetr operates in R environment and was performed as previously described^4^. CellPhoneDB operates in Python environment, and analysis was performed with cpdb_analysis_method.call function using default parameters.

Code in this section is available on Github (https://github.com/FredHutch/Chankhunthod_Sorenson-Graff_et_al_2026).

### Graphics

Some figures were created using BioRender.com.

### Additional statistical analysis

When not specified in prior methods, statistical analyses were carried out using GraphPad Prism (GraphPad Software, La Jolla, CA) in paired t test or unpaired *t* test of cellular phenotypes between two groups.

## Data and code availability

scRNA-seq data used in this article are available through NCBI GEO Series GSE233031. Raw data of living imaging will be available upon request. Codes are accessible through Github: https://github.com/FredHutch/Chankhunthod_Sorenson-Graff_et_al_2026.

## Acknowledgments

This research was supported by NIH R01 HL168110 and R01 HL179317 to B.H., by NIH U24 DK126127 sub-awards (Type A P&F and Type B P&F) and a FHCC/UW/SCH Cancer Consortium Bench-to-Bedside Adult Oncology Award to H.G.Z., and by NIH P30 CA015704 of the Fred Hutch/University of Washington/Seattle Children’s Cancer Consortium, which includes the Flow Cytometry Shared Resource,_RRID:SCR_022613.

## Author contributions

BH and HGZ conceptualized the study, provided research funding, and jointly supervised the study. RSG, JM, YF, and HGZ performed ML analyses. AC, TI, and SD performed experiments, collected data, and analyzed data. RW analyzed data. AC, RSG, BH, and HGZ wrote the manuscript with input from co-authors.

## References

1. Weissman, I.L., and Shizuru, J.A. (2008). The origins of the identification and isolation of hematopoietic stem cells, and their capability to induce donor-specific transplantation tolerance and treat autoimmune diseases. Blood 112, 3543–3553. 10.1182/blood-2008-08-078220.

2. Iwasaki, H., Arai, F., Kubota, Y., Dahl, M., and Suda, T. (2010). Endothelial protein C receptor-expressing hematopoietic stem cells reside in the perisinusoidal niche in fetal liver. Blood 116, 544–553. 10.1182/blood-2009-08-240903.

3. Kim, I., He, S., Yilmaz, O.H., Kiel, M.J., and Morrison, S.J. (2006). Enhanced purification of fetal liver hematopoietic stem cells using SLAM family receptors. Blood 108, 737–744. 10.1182/blood-2005-10-4135.

4. Ishida, T., Mercoli, J., Heck, A.M., Phelps, I., Varnum-Finney, B., Dozono, S., Nourigat-McKay, C., Kraskouskas, K., Wellington, R., Waltner, O., et al. (2025). Differentiation latency and dormancy signatures define fetal liver hematopoietic stem cells at single-cell resolution. Cell Reports 44. 10.1016/j.celrep.2025.116289.

5. Kiel, M.J., Yilmaz, O.H., Iwashita, T., Yilmaz, O.H., Terhorst, C., and Morrison, S.J. (2005). SLAM family receptors distinguish hematopoietic stem and progenitor cells and reveal endothelial niches for stem cells. Cell 121, 1109–1121. 10.1016/j.cell.2005.05.026.

6. Purton, L.E. (2022). Adult murine hematopoietic stem cells and progenitors: an update on their identities, functions, and assays. Exp Hematol 116, 1–14. 10.1016/j.exphem.2022.10.005.

7. Notta, F., Doulatov, S., Laurenti, E., Poeppl, A., Jurisica, I., and Dick, J.E. (2011). Isolation of single human hematopoietic stem cells capable of long-term multilineage engraftment. Science 333, 218–221. 10.1126/science.1201219.

8. Anjos-Afonso, F., Buettner, F., Mian, S.A., Rhys, H., Perez-Lloret, J., Garcia-Albornoz, M., Rastogi, N., Ariza-McNaughton, L., and Bonnet, D. (2022). Single cell analyses identify a highly regenerative and homogenous human CD34+ hematopoietic stem cell population. Nat Commun 13, 2048. 10.1038/s41467-022-29675-w.

9. Varesi, A., Nagree, M.S., Di Biasio, I., Zeng, A.G.X., Shah, S., Zhang, M., Kim, H., Murison, A., Badami, R., Gan, O., et al. (2026). ATP2B1 expression identifies human hematopoietic stem cells with superior repopulation and self-renewal. Proc Natl Acad Sci U S A 123, e2604380123. 10.1073/pnas.2604380123.

10. Fares, I., Chagraoui, J., Lehnertz, B., MacRae, T., Mayotte, N., Tomellini, E., Aubert, L., Roux, P.P., and Sauvageau, G. (2017). EPCR expression marks UM171-expanded CD34(+) cord blood stem cells. Blood 129, 3344–3351. 10.1182/blood-2016-11-750729.

11. Dignum, T., Varnum-Finney, B., Srivatsan, S.R., Dozono, S., Waltner, O., Heck, A.M., Ishida, T., Nourigat-McKay, C., Jackson, D.L., Rafii, S., et al. (2021). Multipotent progenitors and hematopoietic stem cells arise independently from hemogenic endothelium in the mouse embryo. Cell Rep 36, 109675. 10.1016/j.celrep.2021.109675.

12. Rodriguez-Correa, E., Grunschlager, F., Nizharadze, T., Anstee, N., Al-Sabah, J., Kumpost, V., Sedlmeier, A., Li, C., Ball, M., Fotopoulou, F., et al. (2026). A kinetics-based model of haematopoiesis reveals extrinsic regulation of skewed lineage output from stem cells. Nat Cell Biol 28, 1127–1140. 10.1038/s41556-026-01958-0.

13. Tabula Muris, C. (2020). A single-cell transcriptomic atlas characterizes ageing tissues in the mouse. Nature 583, 590–595. 10.1038/s41586-020-2496-1.

14. Zeng, A.G.X., Iacobucci, I., Shah, S., Mitchell, A., Wong, G., Bansal, S., Chen, D., Gao, Q., Kim, H., Kennedy, J.A., et al. (2025). Single-cell Transcriptional Atlas of Human Hematopoiesis Reveals Genetic and Hierarchy-Based Determinants of Aberrant AML Differentiation. Blood Cancer Discov, OF1-OF18. 10.1158/2643-3230.BCD-24-0342.

15. Haas, S., Trumpp, A., and Milsom, M.D. (2018). Causes and Consequences of Hematopoietic Stem Cell Heterogeneity. Cell Stem Cell 22, 627–638. 10.1016/j.stem.2018.04.003.

16. Velten, L., Haas, S.F., Raffel, S., Blaszkiewicz, S., Islam, S., Hennig, B.P., Hirche, C., Lutz, C., Buss, E.C., Nowak, D., et al. (2017). Human haematopoietic stem cell lineage commitment is a continuous process. Nat Cell Biol 19, 271–281. 10.1038/ncb3493.

17. Buggenthin, F., Buettner, F., Hoppe, P.S., Endele, M., Kroiss, M., Strasser, M., Schwarzfischer, M., Loeffler, D., Kokkaliaris, K.D., Hilsenbeck, O., et al. (2017). Prospective identification of hematopoietic lineage choice by deep learning. Nat Methods 14, 403–406. 10.1038/nmeth.4182.

18. Sang-Aram, C., Browaeys, R., Seurinck, R., and Saeys, Y. (2025). Unraveling cell-cell communication with NicheNet by inferring active ligands from transcriptomics data. Nat Protoc 20, 1439–1467. 10.1038/s41596-024-01121-9.

19. Efremova, M., Vento-Tormo, M., Teichmann, S.A., and Vento-Tormo, R. (2020). CellPhoneDB: inferring cell-cell communication from combined expression of multi-subunit ligand-receptor complexes. Nat Protoc 15, 1484–1506. 10.1038/s41596-020-0292-x.

20. Zhang, L., Deng, M., Parthasarathy, R., Wang, L., Mongan, M., Molkentin, J.D., Zheng, Y., and Xia, Y. (2005). MEKK1 transduces activin signals in keratinocytes to induce actin stress fiber formation and migration. Mol Cell Biol 25, 60–65. 10.1128/MCB.25.1.60-65.2005.

21. Murphy, M.J., Wilson, A., and Trumpp, A. (2005). More than just proliferation: Myc function in stem cells. Trends Cell Biol 15, 128–137. 10.1016/j.tcb.2005.01.008.

22. Yin, T., and Li, L. (2006). The stem cell niches in bone. J Clin Invest 116, 1195–1201. 10.1172/JCI28568.

23. Garcia-Prat, L., Kaufmann, K.B., Schneiter, F., Voisin, V., Murison, A., Chen, J., Chan-Seng-Yue, M., Gan, O.I., McLeod, J.L., Smith, S.A., et al. (2021). TFEB-mediated endolysosomal activity controls human hematopoietic stem cell fate. Cell Stem Cell 28, 1838–1850 e1810. 10.1016/j.stem.2021.07.003.

24. Demerdash, Y., Kain, B., Essers, M.A.G., and King, K.Y. (2021). Yin and Yang: The dual effects of interferons on hematopoiesis. Exp Hematol 96, 1–12. 10.1016/j.exphem.2021.02.002.

25. Li, Y., Esain, V., Teng, L., Xu, J., Kwan, W., Frost, I.M., Yzaguirre, A.D., Cai, X., Cortes, M., Maijenburg, M.W., et al. (2014). Inflammatory signaling regulates embryonic hematopoietic stem and progenitor cell production. Genes Dev 28, 2597–2612. 10.1101/gad.253302.114.

26. Christodoulou, C., Spencer, J.A., Yeh, S.A., Turcotte, R., Kokkaliaris, K.D., Panero, R., Ramos, A., Guo, G., Seyedhassantehrani, N., Esipova, T.V., et al. (2020). Live-animal imaging of native haematopoietic stem and progenitor cells. Nature 578, 278–283. 10.1038/s41586-020-1971-z.

27. Wright, D.E., Bowman, E.P., Wagers, A.J., Butcher, E.C., and Weissman, I.L. (2002). Hematopoietic stem cells are uniquely selective in their migratory response to chemokines. J Exp Med 195, 1145–1154. 10.1084/jem.20011284.

28. Aykul, S., and Martinez-Hackert, E. (2016). Transforming Growth Factor-beta Family Ligands Can Function as Antagonists by Competing for Type II Receptor Binding. J Biol Chem 291, 10792–10804. 10.1074/jbc.M115.713487.

29. Hatsell, S.J., Idone, V., Wolken, D.M., Huang, L., Kim, H.J., Wang, L., Wen, X., Nannuru, K.C., Jimenez, J., Xie, L., et al. (2015). ACVR1R206H receptor mutation causes fibrodysplasia ossificans progressiva by imparting responsiveness to activin A. Sci Transl Med 7, 303ra137. 10.1126/scitranslmed.aac4358.

30. Vijaykumar, A., Helbling, P.M., Thelen, F., Fazio, S., Mun, Y., Pereira, A.L., Zielinska, K.A., Buschl, P., Zerjatke, T., Loosli, K., et al. (2026). Tissue-scale mapping reveals a central role of hepatoblasts in the regulation of fetal liver hematopoiesis and stem cell maintenance. Dev Cell 61, 901–918 e904. 10.1016/j.devcel.2026.01.011.

31. Loeffler, D., Wehling, A., Schneiter, F., Zhang, Y., Muller-Botticher, N., Hoppe, P.S., Hilsenbeck, O., Kokkaliaris, K.D., Endele, M., and Schroeder, T. (2019). Asymmetric lysosome inheritance predicts activation of haematopoietic stem cells. Nature 573, 426–429. 10.1038/s41586-019-1531-6.

32. Arif, T., Qiu, J., Khademian, H., Lohithakshan, A., Menon, A., Menon, V., Slavinsky, M., Batignes, M., Lin, M., Sebra, R., et al. (2025). Reversing lysosomal dysfunction restores youthful state in aged hematopoietic stem cells. Cell Stem Cell 32, 1904–1922 e1907. 10.1016/j.stem.2025.10.012.

33. Wolf, F.A., Angerer, P., and Theis, F.J. (2018). SCANPY: large-scale single-cell gene expression data analysis. Genome Biol 19, 15. 10.1186/s13059-017-1382-0.

34. Chen, J., Bardes, E.E., Aronow, B.J., and Jegga, A.G. (2009). ToppGene Suite for gene list enrichment analysis and candidate gene prioritization. Nucleic Acids Res 37, W305–311. 10.1093/nar/gkp427.

