## Supplementary material for "Single-Cell Behavioromics Enables Fate Prediction of Individual Hematopoietic Stem Cells": Suppl

#### Supplementary Figure Legends

##### Supplementary Figure 1.

**(A)** Flow cytometry gating strategy for immunophenotyping of single HSPC-derived colonies following 14 days of co-culture with FL-EC. **(B)** Representative image sequences of single HSPCs from Cluster 1 and Cluster 3 from Figure 1E. Images are in greyscale showing the pixel intensity of CellTrace Far Red signal. Blue circles mark the cell locations in zoomed-in frames of the whole-well images. t: time elapsed. **(C)** Stochasity of cell movements was calculated as Shannon Entropy of the directions of cell location changes between every two frames. This shows that the movements of both serial-LT-HSCs and other HSPCs are stochastic and not uni-directional.

##### Supplementary Figure 2.

(**A**) Significantly enriched GO terms (biological process) among downregulated genes in serial-LT-HSCs compared to other HSPCs. GOs are color-coded by their functional categories. **(B)** Flow cytometry density plots showing representative examples of defined immunophenotypes of colonies derived from single serial-LT-HSCs, ST-engrafting progenitors, and non-engrafting progenitors. Only SCA1, EPCR, and ESAM are shown. $CD45^{+}GR1^{-}F4/80^{-}$ phenotype is shared across all HSPCs in this study.

### Supplementary Figure 1

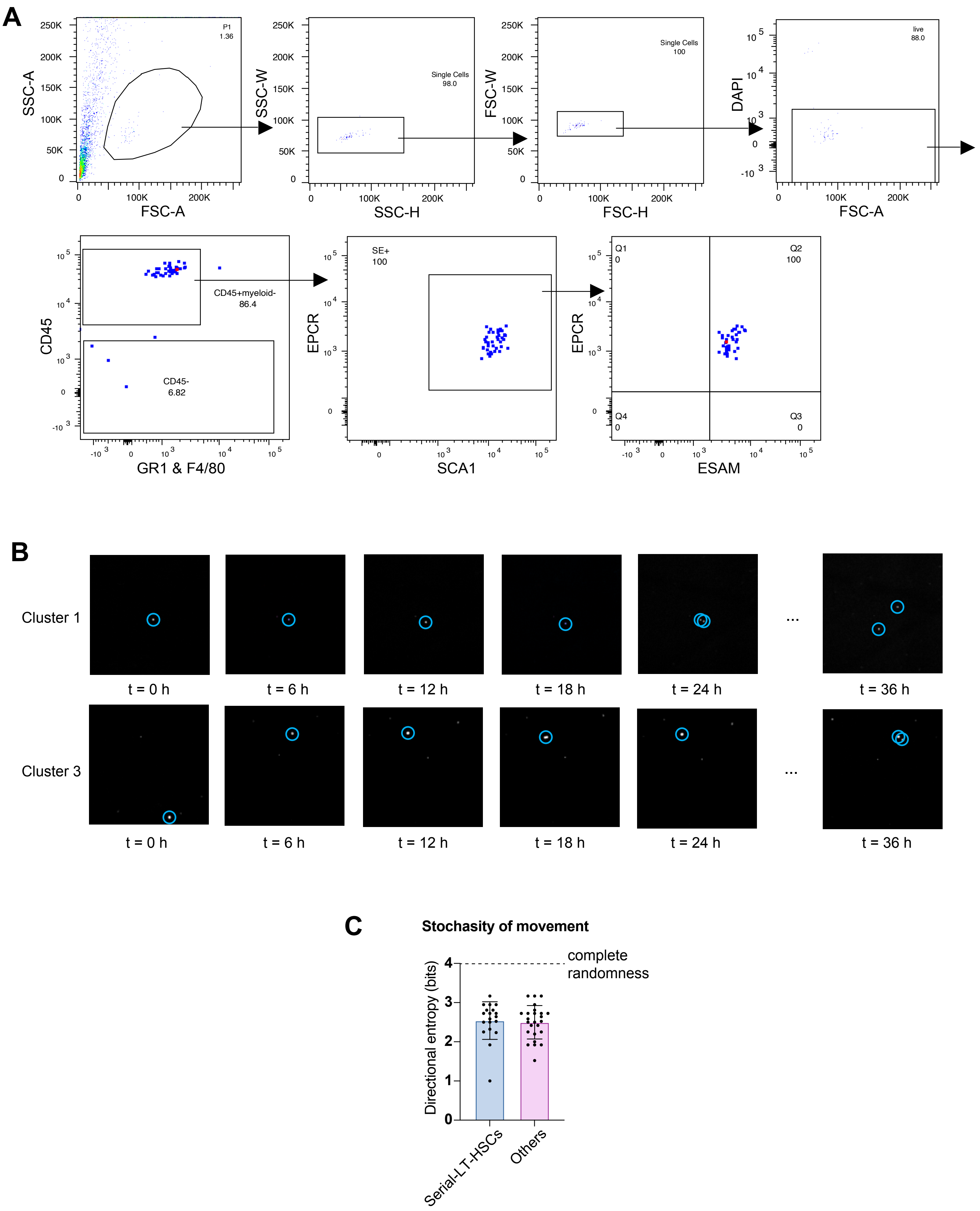

### Supplementary Figure 2

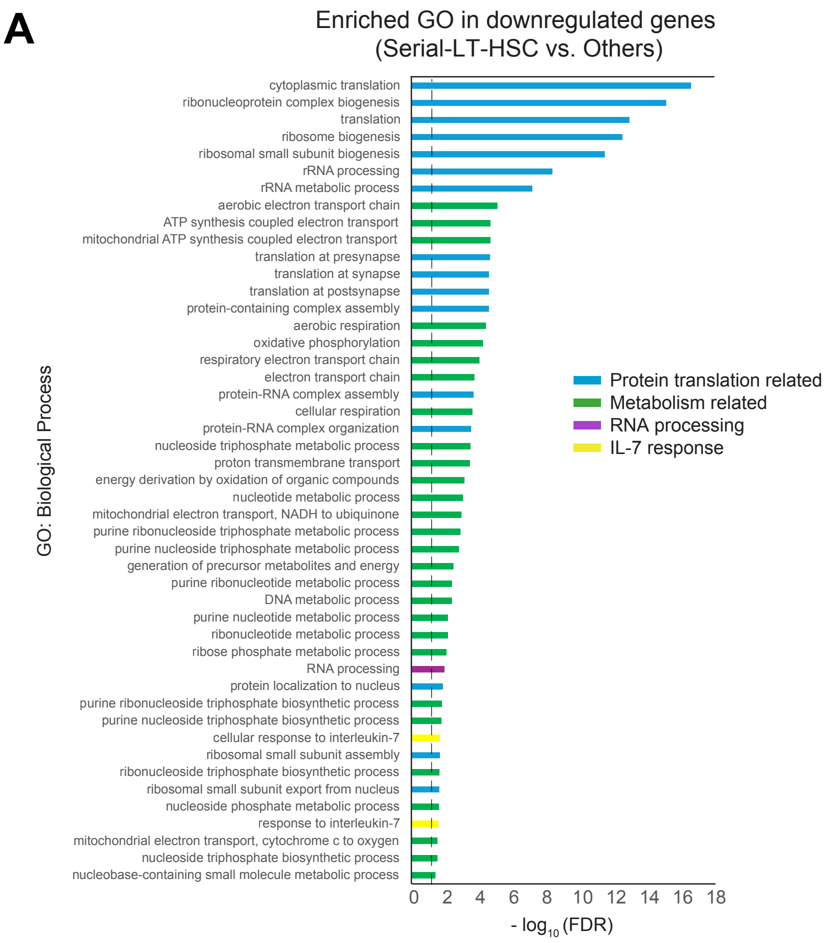


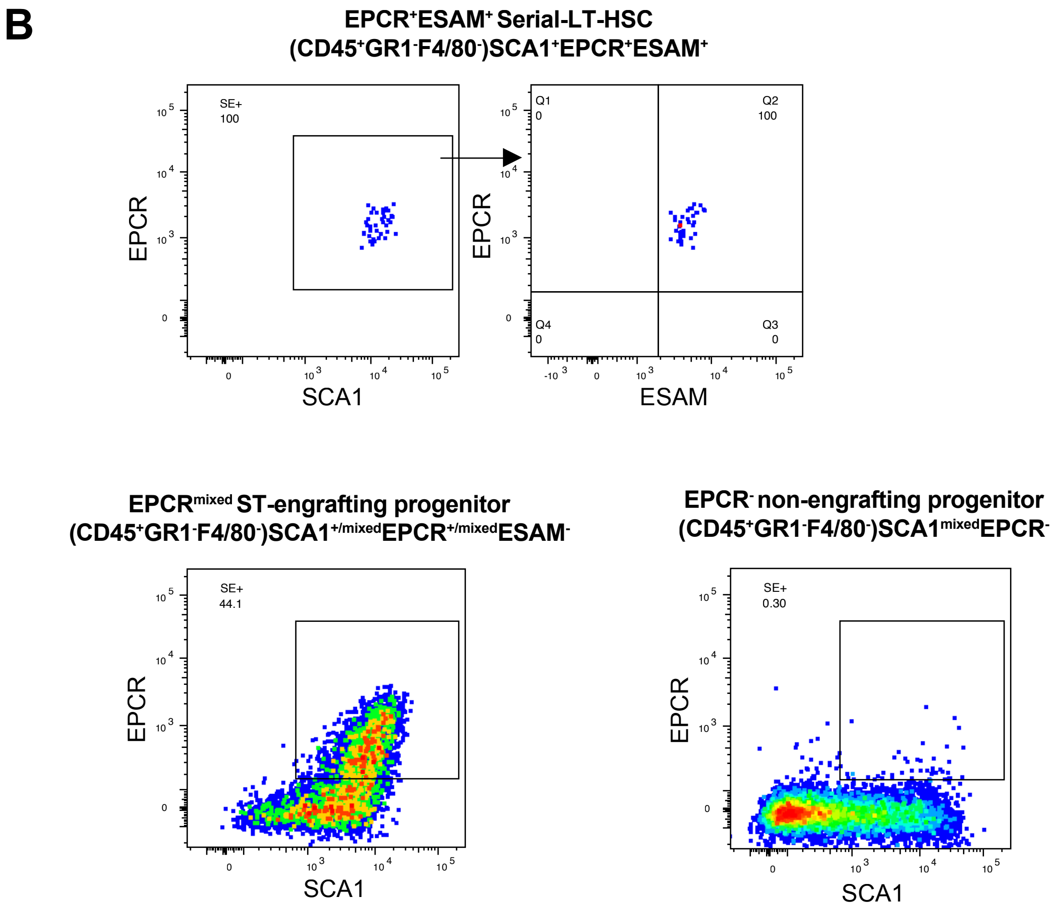
